# Complex deletion and proximal reinsertion of a 150bp regulatory sequence in the mouse Csf1r promoter mediated by CRISPR-Cas9

**DOI:** 10.1101/872200

**Authors:** Kim M. Summers, Clare Pridans, Evi Wollscheid-Lengeling, Kathleen Grabert, Antony Adamson, Neil E. Humphreys, Katharine M. Irvine, David A. Hume

## Abstract

This paper describes a deletion/reinsertion event encountered in a genome-editing project using CRISPR-Cas9. The objective was to delete a 150bp enhancer region in the mouse *Csf1r* locus using a pair of guides and a homology-dependent repair (HDR) template. The editing was successful in generating a founder pup with the anticipated precise deletion. However, the deleted fragment and a duplicated copy of part of the HDR template was reinserted around 50bp downstream. The reinsertion event was recognised because the PCR primer site used in genotyping was duplicated, so that there were three PCR products in a heterozygous animal and two in a homozygote. The event we describe is more subtle and more difficult to detect than large-scale rearrangements reported by others. We suggest that any genomic deletion mediated by CRISPR-Cas9 needs to be confirmed by assessing the copy number in the genome.

## Introduction

The development of the CRISPR-Cas9 technology for genome editing (1, 2) revolutionised functional genomics in the mouse. DsDNA breaks or deletions directed by individual or pairs of guide RNAs can be repaired by non-homologous end-joining (NHEJ) but the extent and direction of such deletions is unpredictable (3). Further development of the genome-editing technology to favour homology-directed repair (HDR) of dsDNA breaks generated by CRISPR-Cas9 has facilitated the generation of point mutations and larger insertions and deletions (e.g reporter genes, conditional alleles etc) in the mouse genome (e.g. (4–7)). despite the many advances, off-target events and unpredictable on-target effects including deletions and rearrangements remain a concern (8, 9).

Our research is focussed on transcriptional regulation in the macrophage lineage. Amongst the many applications of CRISPR-Cas9 technology has been the dissection of the functions of individual enhancers in myeloid-specific transcription factor loci (10–13). We recently deleted a highly-conserved intronic enhancer in the *Csf1r* locus using CRISPR-Cas9 (14). Homozygous mutant mice were *Csf1r* hypomorphs, lacking *Csf1r* expression in monocytes and a subset of *Csf1r-*dependent tissue macrophage populations. These findings led us to examine the functions of other regulatory elements in the *Csf1r* locus. One such element is a 150bp region upstream of the transcription start site that is required for maximal expression of reporter genes in transgenic mice (15) and includes a promoter that is used by the placental trophoblast. We therefore devised a strategy to delete this region in the mouse germ line using homology-direct repair. The strategy was successful at the intended target site, but subsequent analysis revealed that the deleted segment was reinserted immediately downstream. This outcome highlights the need for caution in analysis of the results of genome editing.

## Methods

The generation of CRISPR-Cas9 deletions by pronuclear injection of mouse embryos was carried out essentially as described previously (16, 17). The initial strategy shown in Figure 1 and involved the coinjection of two guides, a ssDNA homology-directed repair (HDR) template spanning the desired deletion and recombinant Cas9. Two separate rounds of injections used different guides at the 5’ end of the desired deletion.

**Figure 1.**
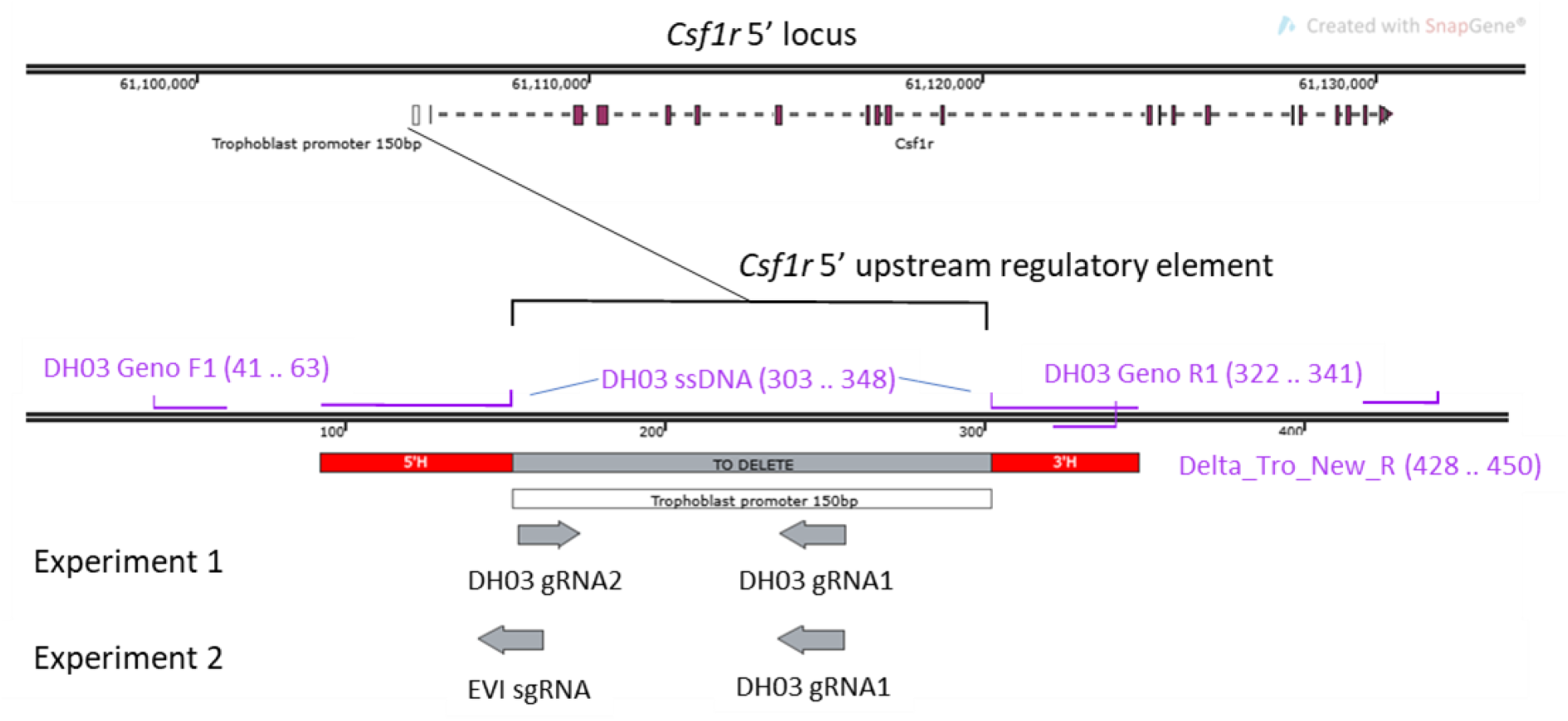
Strategy for deletion of Csf1r 150 bp upstream regulatory region. Two sets of guides were designed. Experiment 1 used DH03 gRNA1 and DH03 gRNA2; experiment 2 used DH03 gRNA1 and EVI sgRNA. DH03 Geno F1 and DH03 Geno R1 (shown in purple) were used for the initial genotyping. Delta_Tro_New_R was used with DH03 GenoF1 to sequence beyond the target region. Inner purple sequences show the oligonucleotide used for homology directed repair, also shown in red boxes; grey box shows the region to be deleted.

Guides were designed using the Sanger website using stringent criteria for off target predictions (guides with mismatch (MM) of 0, 1 or 2 for elsewhere in the genome were discounted; MM3 were tolerated if predicted off targets were NOT exonic). A separate guide RNA was designed based on *in vitro* data from Evi. An Alt-R crRNA (IDT) oligo for the chosen guide was ordered and resuspended in sterile, RNase free injection buffer (TrisHCl 1mM, pH 7.5, EDTA 0.1mM) and annealed with tracrRNA (IDT) by combining 2.5ug crRNA with 5ug tracrRNA and heating to 95oC. The mix was allowed to slowly cool to room temperature. After annealing the complex an equimolar amount was mixed with 1000ng Cas9 recombinant protein (NEB; final conc 20ng/ul) and incubated at RT for 15’, before adding Cas9 mRNA (final conc; 20ng/ul) and the ssDNA PAGE purified repair template (IDT; final conc 50ng/ul) in a total injection buffer volume of 50ul. The injection mix was centrifuged for 10’ at RT and the top 40ul removed to another tube for injection.

Two days of microinjections were performed using the AltR crRNA:tracrRNA:Cas9 complex (20ng/ul; 20ng/ul; 20ng/ul respectively), Cas9 mRNA (20ng/ul), and ssDNA HDR template (50ng/ul). This mix was microinjected into the pronuclei of one-day single cell mouse embryos. Zygotes were cultured overnight and the resulting 2 cell embryos surgically implanted into the oviduct of day 0.5 post-coitum pseudopregnant mice.

After birth and weaning genomic DNA was extracted with the Sigma redextract-n-amp tissue PCR kit and used to genotype pups. PCR was performed using primers DH03 Geno F1 tgctataggaggctcattactgt and DH03 Geno R1 caccacacaccagaggaaga (Figure 1) which yielded a 301bp band on WT template genomic DNA. Amplicons were run on Qiaxcel (Qiagen) and pups demonstrating reduction in predicted band size were taken forward for sequencing. Using high fidelity polymerases (Phusion, NEB), the amplicons were subcloned in pCRBlunt (Invitrogen) and Sanger sequenced using M13 F and R primers. Sequences were aligned to WT sequence in Snapgene (GSL Biotech) to confirm exact deletion.

One F1 male pup was mated to a wild type female and the progeny genotyped as described above. The amplified bands were eluted from the gel using the Qiagen gel extraction kit. Eluted bands were sent for chain termination sequencing using the genotyping primers (Figure 1). Subsequently, an additional reverse primer (Delta_Tro_new_R; agatgaacacgttcctgtctgtt) was used with DH03 GenoF1, to amplify and sequence beyond the target region.

## Results and Discussion

Figure 2A shows the PCR genotyping of individual pups from the first-round targeting. Two pups (9 and 10) showed evidence of a truncated band amplified using primers flanking the deleted region. However, the band in each case was larger than anticipated for a full 150bp deletion. In both cases, there was evidence of an amplified band that was larger than the predicted wild type band. These bands were considered to result from annealing between the deleted fragment and the WT fragment forming a heteroduplex in amplified DNA from heterozygous animals and were not characterised further. The smaller bands in Pups 9 and 10 were isolated and sequenced. The results (Figure 2B) indicated partial deletions of the target region. The outcome suggested that the deletions arose from a single site of dsDNA cleavage at gRNA1 and did not involve HDR. Larger bands were also observed in 3 other pups (3, 5 and 8).

**Figure 2.**
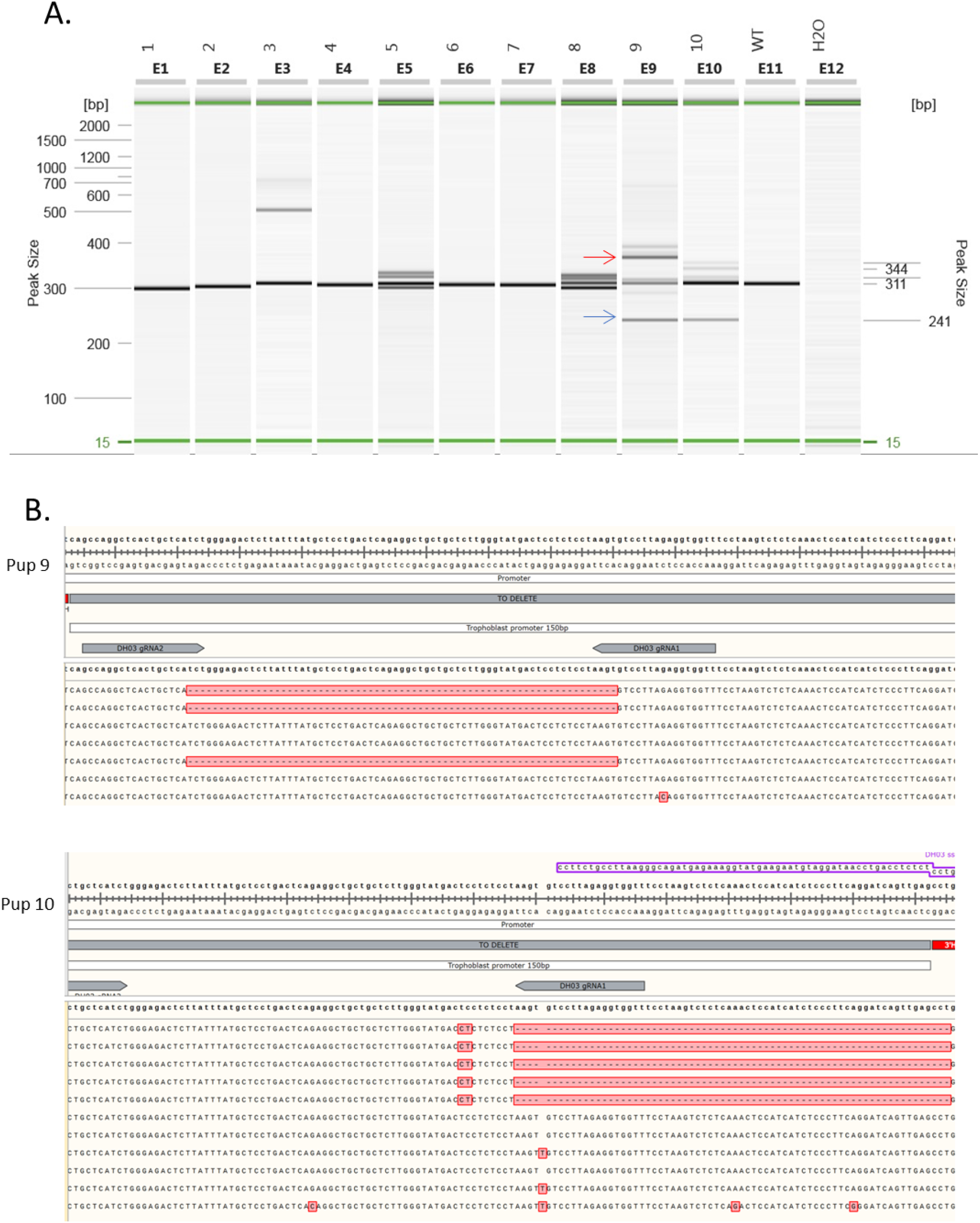
Results from first round targeting. A. PCR genotyping of pups from first round targeting. Primers DH03 Geno F1 and DH03 Geno F2 were used. Wild type band is at 311 bp, blue arrow shows the smaller band in pups 9 and 10; red arrow shows the larger putative heteroduplex band. B. Sequencing of *Csf1r* in Pups 9 and 10 from first round targeting.

Accordingly, in a second round of targeting, we designed an alternative 5’ guide RNA and tested its ability to generate dsDNA cleavage and deletions in transfected RAW264 cells ((14); not shown). Using the alternative guide, and the same PCR genotyping of the progeny generated following pronuclear injection, we identified 1 pup with the expected deletion (Pup 16) and another (Pup 12) with a smaller deletion (Figure 3A). In both cases, a PCR product of around 400 bp, larger than the expected wild-type product, was also observed. This was again attributed to heteroduplex formation between products from the two alleles. The smaller band of the anticipated size for a 150bp deletion from Pup16 was cloned and sequenced and found to contain the precise deletion anticipated following successful HDR (Figure 3B).

**Figure 3.**
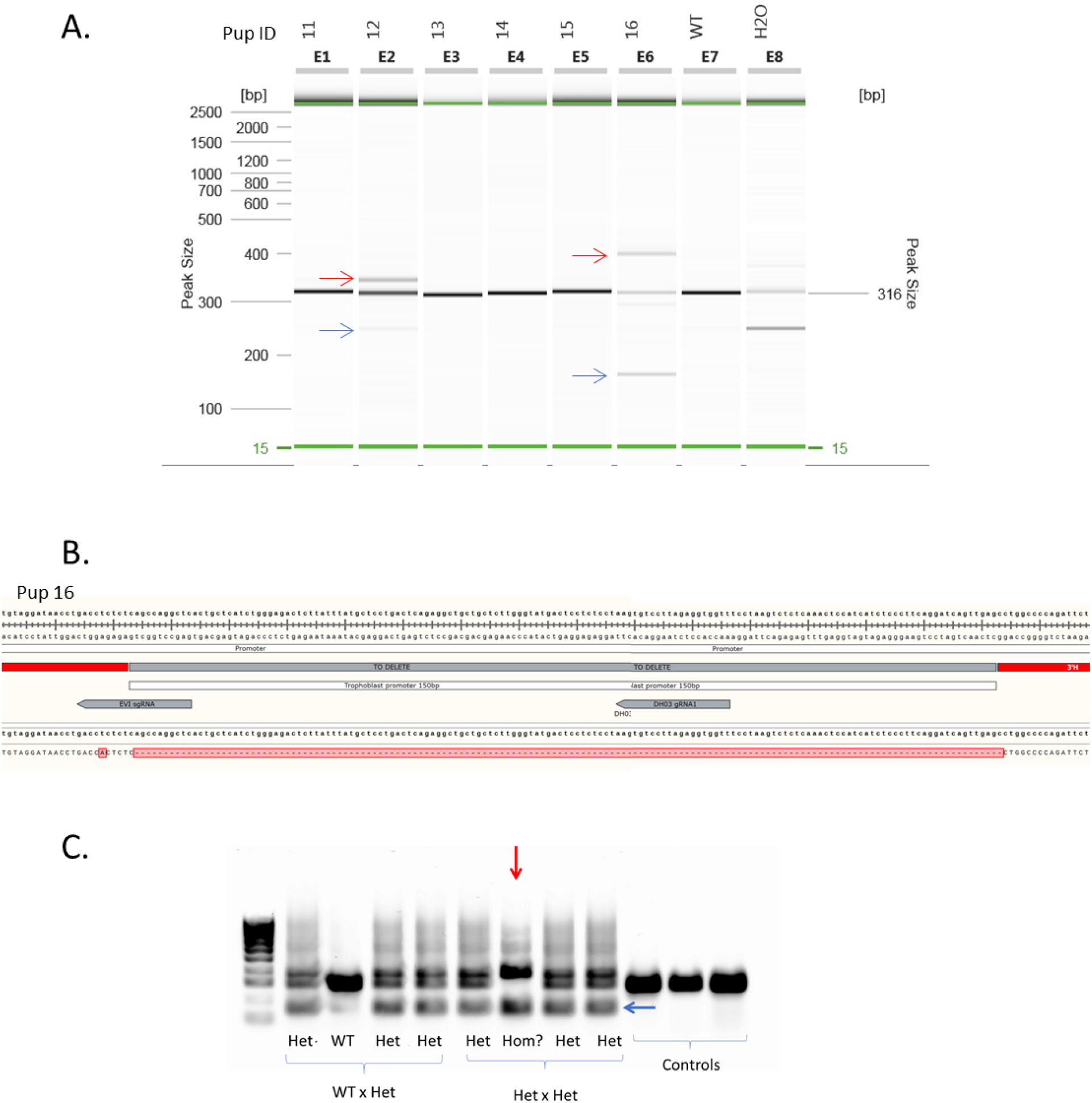
Results from second round targeting. A. PCR genotyping of pups from second round targeting. Primers DH03 Geno F1 and DH03 Geno F2 were used. Wild type band is at 316 bp, blue arrows show the smaller bands in pups 12 and 16; red arrows show the larger putative heteroduplex bands. B. Sequencing of *Csf1r* in Pup 16 from second round targeting. C. PCR genotyping of F2 and back cross progeny. Blue arrow shows the smaller, deletion band. Four offspring are from a heterozygous/wild type mating (WT × Het); four are from a heterozygous/heterozygous mating (Het × Het). Controls were from a different mouse strain. Putative genotypes are shown above the cross. Red arrow shows the sample with the smaller and larger bands but without the wild type band, suggesting that it is homozygous for the deleted region. Molecular weight ladder is BioLine HyperLadder 100bp with markers at 100 bp intervals.

The male Pup 16 was mated to a wild-type mouse and heterozygous progeny from that mating were backcrossed to generate homozygotes. In the F2 generation, we obtained one offspring in which the ~400bp larger and 151bp smaller bands were both present and the 301 bp wild-type band was absent (Figure 3C), indicating that the larger band could not be a heteroduplex. To determine the nature of the two bands, both were isolated and sequenced. Comparison of the two sequences revealed that the mutant *Csf1r* allele contained the 150bp deletion at the anticipated location as well as a downstream insertion that contained the entire 150bp deletion plus a partial duplication of the 3’ side of the HDR template. The duplicated HDR sequence contained the PCR primer site used in genotyping, explaining the generation of two bands (Figure 4). The insertion does not disrupt conserved elements of the proximal promoter. It is unlikely to alter transcription, but in any case, any phenotype associated with the mutant allele would be uninterpretable.

**Figure 4.**
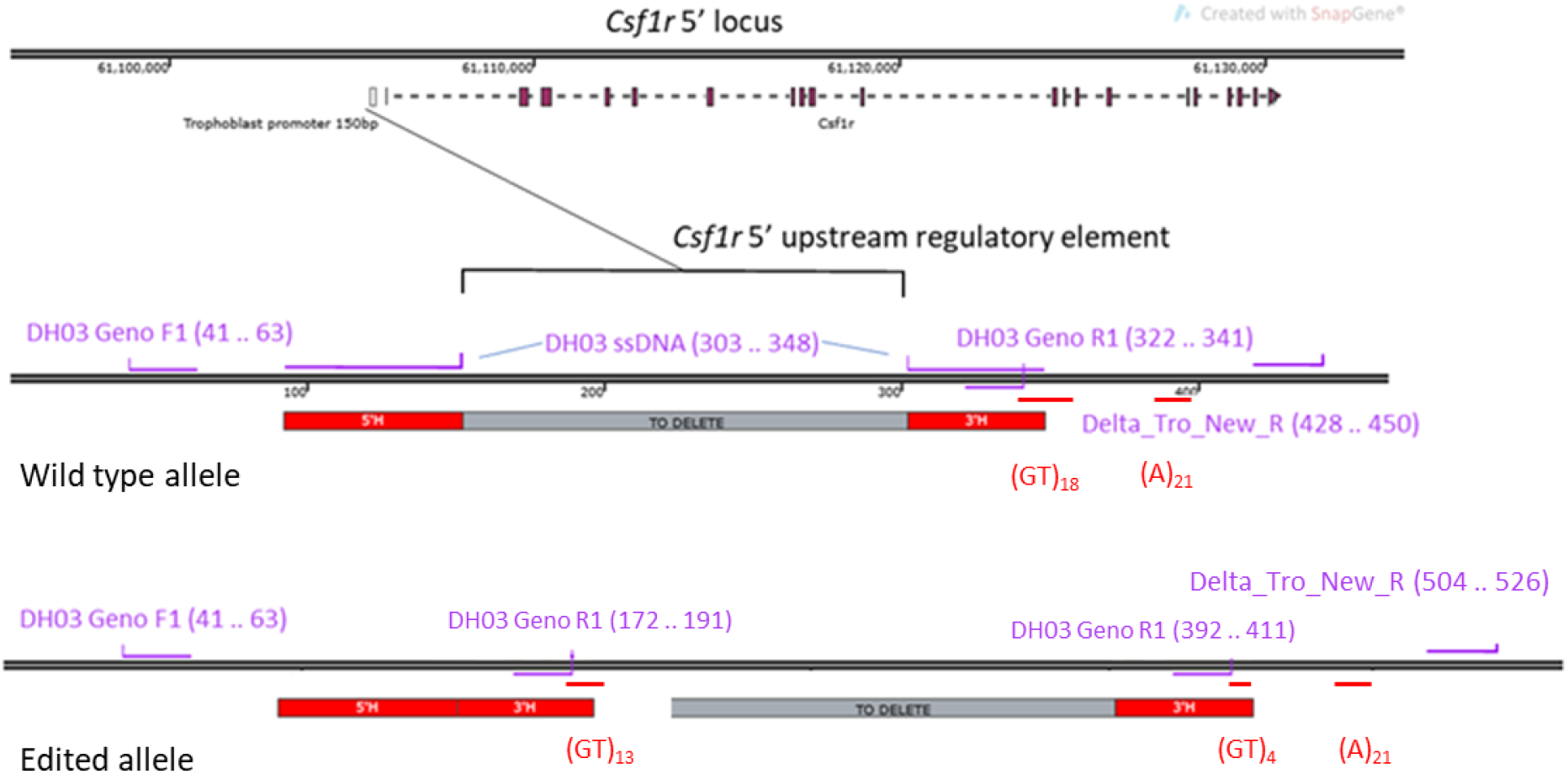
Proposed structure of the edited allele. The two copies of the reverse primer DH03 GenoR1 lie 200 bases apart; the second reverse primer lies about 100 bases downstream. Primers are shown in purple; repeat sequences that make editing and sequencing problematic are shown in red.

The sequence of the *Csf1r* locus does not reveal an obvious alternative binding site for the guide RNAs, although there is growing evidence of off-target cleavage by Cas9 (8) and cleavage of sites adjacent to the targeted location is likely (9). The structure of the target locus suggests a possible mechanism or at least a sequence to avoid in CRISPR-Cas9 targeting. There is an extended GT/CA repeat (25 copies) immediately 3’ of the target sequence and partly overlapping the 3’ arm of the HDR template (Figure 4). GT repeat sequences are associated with Z-DNA structure and recombination hot spots in the genome (18, 19). The derived sequence following the reinsertion event contains a partial deletion of the GT repeat (Figure 4) that might have arisen through recombination. A recent study demonstrated that extended GT/CA repeats can also bind Ku70/80 (19) which is required for non-homologous end-joining (18). In addition, the left hand guide contains the sequence AGAGA which is found at the breakpoint of the insertion and may have facilitated this process.

Complex large-scale deletions and insertions arising from CRISPR-Cas9 editing have been reported by others working in cell lines and transgenic mice (3, 5, 9). A recent study reported a so-called bystander mutation arising from deletion of an enhancer in the mouse *Il2ra* gene, a large tandem duplication of adjacent sequences (20). In our case, the intended deletion is relatively small (typical of enhancer deletions). The re-insertion is much more subtle and might easily have been missed if it had not also duplicated the 3’ end of the HDR template containing the primer sites used for PCR genotyping. Had this not occurred, we would have detected a single PCR product with the precise deletion required and progressed to conclude that the deletion had no effect on transcription of *Csf1r.* Based upon the analysis of Pup 16, we strongly suspect that larger bands seen in other founders also reflect insertions/duplications. To avoid this kind of artefact in the future, we would advocate testing founders for reinsertion events by Southern Blotting using the excised fragment as a probe rather than relying solely on PCR analysis.

In overview, this project highlights an unusual and specific form of bystander mutagenesis of a target locus by CRISPR-Cas9 that may actually have been promoted by the use of HDR, or indeed a combination of repair pathways. Whilst in our experience these atypical repair outcomes are uncommon, reporting of these events is vital to build a more detailed overview of potential gene editing outcomes. This is especially relevant in the context of using gene editing in the clinic.

